# IL-2Rα KO mice exhibit maternal microchimerism and reveal nuclear localization of IL-2Rα in lymphoid and non-lymphoid cells

**DOI:** 10.1101/2023.11.03.565571

**Authors:** Victoria A. Wong, Kristie N. Dinh, Guangchun Chen, Lucile E. Wrenshall

## Abstract

IL-2Rα KO mice have been instrumental to discovering the immunoregulatory properties of IL-2Rα. While initially thought of only as a stimulatory cytokine, IL-2 and IL-2Rα knock out (KO) mice revealed that this cytokine-receptor system controls immune responses through restimulation-induced cell death and by promoting the survival of T regulatory cells. Although described mostly in the context of lymphocytes, recent studies by our laboratory showed that IL-2R is expressed in smooth muscle cells. Given this finding, we sought to use IL-2Rα knock mice to determine the function of this receptor in vascular smooth muscle cells. Surprisingly, we found that IL-2Rα knock out vascular smooth muscle cells had detectable IL-2Rα. Further studies suggested that the source of IL-2Rα protein was likely maternal heterozygous cells present in KO offspring due to maternal microchimerism. Because the KO was generated by using a neomycin resistance gene insert, we treated cells with G418 and were able to eliminate the majority of IL-2Rα expressing cells. This elimination revealed that IL-2Rα KO vascular smooth muscle cells exhibited increased proliferation, decreased size, and hypodiploid DNA content when compared to wildtype cells. Our findings suggest that the phenotype of complete IL-2Rα loss is more severe than demonstrated by IL-2Rα KO mice, and that IL-2Rα plays a here-to-fore unrecognized role in regulating cell proliferation in non-lymphoid cells.

## Introduction

IL-2Rα, or CD25, is a key subunit of the tripartite IL-2 receptor. Expression of IL-2Rα distinguishes activated from naïve T cells, and high expression serves as a marker for T regulatory cells. The contribution of IL-2Rα to T regulatory cell survival and prevention of autoimmunity has been demonstrated through IL-2Rα knock out (KO) mice, which develop splenomegaly, lymphadenopathy, anemia, and inflammatory bowel disease (1,2). IL-2Rα KO mice have been instrumental in determining the impact of IL-2Rα on the immune system (1,3–5).

Although the IL-2R is best known for its expression on lymphocytes, there are limited reports demonstrating the expression of IL-2R on other cell types, particularly dendritic cells (6–8). Our laboratory recently reported that vascular smooth muscle cells (VSMC) express all 3 subunits of the IL-2R (9). We found that expression of the IL-2Rα differed with VSMC phenotype, with proliferating VSMC expressing lower levels of IL-2Rα than differentiated VSMC.

To help define the function of IL-2Rα in VSMC, we sought to study cell properties in the absence of IL-2Rα. To this end, we isolated VSMC from IL-2Rα wildtype (WT) and KO mice. Surprisingly, we found that IL-2Rα protein was present, mainly in the nucleus, of both WT and KO VSMC. IL-2Rα was detected in IL-2Rα KO VSMC and splenocytes by immunofluorescence, flow cytometry, and Western blot analysis. Low levels of WT *IL2RA* were detected in genomic DNA extracted from multiple IL-2Rα KO tissues. Our studies suggest that the source of *IL2RA* DNA, and in turn IL-2Rα protein, is maternal microchimerism. The magnitude and pervasive nature of the IL-2Rα protein in KO mice is likely due to our finding that IL-2Rα protein has a surprisingly long half-life, and that IL-2Rα protein can be transferred between cells.

## Materials and Methods

### Materials and animals

Animal studies were approved by the Institutional Animal Care and Use Committee at Wright State University and conformed to the Guide for the Care and Use of Laboratory Animals (NIH Publication, 8^th^ edition). Heterozygous breeding pairs of IL-2Rα null (B6;129S4-*IL-2ra*^tm1Dw^/J) mice were purchased from The Jackson Laboratory (Bar Harbor, ME, USA) for the establishment of breeding colonies (1).

Antibodies recognizing IL-2Rα were from the following sources: rabbit anti-human polyclonal anti-IL-2Rα antibodies were from Bioss Antibodies, Inc. (Woburn, MA), BosterBio (Pleasanton, CA), and LSBio (Seattle, WA). Mouse anti-human and anti-mouse IL-2Rα, clone 1B5D12, was from Genetex (Irvine, CA). Rabbit anti-phospho-CD25 (ser268) was from Sigma (St. Louis, MO). Rat anti-mouse CD16/32, rat anti-mouse CD4, and rat anti-mouse IL-2Rα (clone PC61) were from Biolegend (San Diego, CA). Purified mouse IL-2Rα was from SinoBiological (Wayne, PA). Antibodies used for SMC identification were as follows: rabbit anti-human smooth muscle cell alpha actin was from Novus (Littleton, CO). Sheep anti-transgelin (multiple species) was from R&D Systems (Minneapolis, MN). Rabbit anti-human h-caldesmon was from Proteintech (Rosemont, IL). Benzonase® was from Minnebio (St. Paul, MN). DMEM was from ThermoFisher (Waltham, MA).

### VSMC culture

Murine VSMC were isolated from thoraco-abdominal aortas by enzymatic digestion based on a protocol from Kwartler, et al (10). Aortas were dissected free of adipose tissue *in situ*, removed and rinsed three times in sterile Hanks’ balanced salt solution, then digested at 37°C for 16-18h with a mixture of 0.1 mg/ml collagenase (Sigma), 25 μg/ml trypsin inhibitor (Sigma) and 18 μg/ml elastase (Worthington Biochemical Corp., Lakewood, NJ). Isolated smooth muscle cells and any remaining tissues were then washed and cultured in DMEM with 10% FBS at 37°C and 5% CO_2_. Tissue debris was removed the following day.

For serum free conditions, VSMC were cultured in DMEM supplemented with insulin, transferrin, and selenium (ITS; Sigma) (11). Identification of cells as VSMC was confirmed through expression of smoothelin (passage 1-2 only), smooth muscle cell α-actin, transgelin, and h-caldesmon.

Human VSMC were isolated from pieces of aorta using the explant technique (9). Tissues were washed with PBS and all adipose was removed. The artery was then cut into small (approximately 5 mm x 5 mm) pieces and 2-3 pieces per well were placed lumen side down into 6 well tissue culture plates. Pieces were allowed to adhere briefly, then smooth muscle cell media (ScienCell, Carlsbad, CA) supplemented with 10% FBS was carefully added to the wells so as not to dislodge the pieces of aorta. The tissue was maintained at 37°C and SMCs were harvested in 3 - 4 weeks. VSMC were then passaged every 7 - 8 days and used after 1-2 passages.

### PCR and qPCR

Offspring of heterozygous breeding pairs of IL-2Rα null (B6;129S4-*IL-2ra*^tm1Dw^/J) mice were genotyped per Jax mice touchdown protocol (12). Briefly, DNA was extracted from tail clippings and amplified using the recommended primers and touchdown protocol. Primers were as follows: (WT forward CTGTGTCTGTATGACCCACC, WT reverse CAGGAGTTTCCTAAGCAACG; mutant forward CTTGGGTGGAGAGGCTATTC, mutant reverse AGGTGAGATGACAGGAGATC). The WT primer sequences are contained within exon 2.

Quantitative real time PCR was performed, using the above primers, with 15 ng genomic DNA input in triplicate on a 7500 fast real-time PCR system (Applied Biosystems, ThermoFisher Scientific) using PowerUp™ SYBR™ Green Master Mix kit (Applied Biosystems) according to the manufacturer’s instructions. The cycle profile was 2 minutes at 50 °C, 2 minutes at 95 °C, 40 cycles of 3 seconds at 95 °C and 30 seconds at 60 °C. GAPDH was used as an endogenous control. RT-minus and no template controls (NTC) were included for negative control. Data were analyzed using 7500 software v2.3 (Applied Biosystems).

### Flow cytometry

Splenocyte cell suspensions were prepared by mechanical disruption of the spleen followed by filtration to remove large pieces of noncellular debris. The following incubations were all performed on ice. Splenocytes were first incubated with anti-CD16 (TruStain, FcX) for 15 min, then washed and stained with the indicated anti-IL-2Rα antibodies for 30 minutes. Following fixation in 2% paraformaldehyde, intracellular staining was performed by first permeabilizing cells via suspension in ice cold methanol for 30 minutes. Cells were then washed twice in PBS with 2% BSA and stained with anti-IL-2Rα antibodies for 30 minutes. Clone PC 61.5 and anti-IL-2Rα from Genetex were directly conjugated to Cy-5 fluorochromes and anti-IL-2Rα from Boster was used with a Cy5 labeled secondary antibody. A subset of cells was treated with Benzonase® 250U/10^6^ cells for 50 minutes at 37°C prior to staining. Data were acquired on a Accuri C6 flow cytometer (Bectin Dickinson, Franklin Lakes, NJ) and analyzed using FCS Express 7 (Pasadena, CA).

### Microscopy

VSMC were cultured in chamber slides and fixed in ice cold methanol with 2% acetic acid. Fixed VSMC were then washed with PBS and blocked for 1h using Odyssey blocking solution (LI-COR, Lincoln, NE). Blocking solution was removed by washing, and primary antibodies were diluted in TBST (10 mM Tris, pH 8.0, 150 mM NaCl, 0.5% Tween 20) and applied to cells overnight at 4°C. The primary antibody was removed by washing, and the appropriate fluor-conjugated secondary was applied for 1h. Cells were imaged on an EVOS epifluorescent microscope (ThermoFisher) or on a Cytation imaging plate reader (Biotek, Winooski, VT). Images from the Cytation were processed using the flow cytometry software FCS Express 7.

### Western blot analysis

Whole cell lysates from primary VSMCs and Jurkat T cells (ATCC, Manassas, VA) were prepared by lysing and sonicating pelleted cells in radio immuno-precipitation (RIPA) buffer (150 mM NaCl, 25 mM Tris-HCl pH 7.6, 1% Nonidet P40, 1.0 % sodium deoxycholate, 0.1 % SDS). Splenocytes were separated into membrane, cytoplasmic, and nuclear fractions per manufacturer’s instructions (Cell Fractionation Kit, Cell Signaling Technology, Danvers, MA). Extracts were then separated by SDS-PAGE, using 30 μg protein measured by bicinchoninic acid assay (Pierce, Thermo-Fisher), and transferred to a polyvinylidene difluoride membrane. Some blots, as indicated in figure legends, were assessed for total protein using either stain-free gels (Bio-Rad, Hercules, CA) or the Revert™ 700 Total Protein Stain (LI-COR) prior to blocking. After incubation with 5% nonfat milk in TBST for 60 min, the membrane was incubated with antibodies against IL-2Rα at 4 °C for 18 h. Membranes were washed three times for 10 min and incubated with a 1:10,000 dilution of horseradish peroxidase-conjugated anti-rabbit antibodies (Jackson ImmunoResearch, West Grove, PA) at room temperature for 2 hrs. Blots were washed with TBST three times and developed with the ECL system (Luminata Crescendo, Millipore Sigma) according to the manufacturer’s instructions.

### Protein half-life

The half-life of IL-2Rα was measured using a protocol designed by Morey, et al (13) in which a traditional pulse-chase experiment was performed using L-azidohomoalanine (AHA), a methionine analog, as the pulsed label. Incorporation of AHA into nascent proteins was detected by a click chemistry based, copper-free strain-promoted alkyne-azide cycloaddition reaction in which a fluorescent cyclooctyne probe (dibenzocyclooctyne-488) bound to incorporated AHA (14,15).

Specifically, VSMC were washed then cultured in methionine free DMEM with 5% dialyzed FBS containing 50 μM L-azidohomoalanine (AHA) for 48 hours (duration determined by pilot experiments) then cells were returned to DMEM with 10% FBS. VSMC were pelleted at indicated time intervals following this initial pulse and then processed together. Because IL-2Rα is primarily localized to the nucleus in VSMC, we focused on extracting nuclear proteins. Nuclear fractions were isolated following the protocol outlined by Senichkin, et al (16). Briefly, pelleted cells were incubated in a hypotonic buffer (20 mM Tris-HCl (pH 7.4), 10 mM KCl, 2 mM MgCl_2_, 1 mM EGTA, 0.5 mM DTT, 0.5 mM PMSF) for 3 minutes, followed by addition of 0.1% NP-40 (final concentration) for an additional 3 minutes. Following centrifugation, the pellet was incubated in isotonic buffer (20 mM Tris-HCl (pH 7.4), 150 mM KCl, 2 mM MgCl_2_, 1 mM EGTA, 0.5 mM DTT, 0.5 mM PMSF) plus 0.1% NP-40 for another 3 minutes, then centrifuged. The resulting nuclear pellet was resuspended in 100 μl Benzonase^®^, 100U for 10 minutes at 37°C. RIPA buffer, 400 μl, was then added for 20 minutes at room temperature (RT). Following centrifugation, the supernatant (nuclear soluble fraction) was saved and the insoluble pellet, if present, was washed and resuspended in 100 μl Benzonase^®^ 100 U. Following a 10-minute digestion at 37°C, the digested lysate was combined with the nuclear soluble fraction for immunoprecipitation, labeling with dibenzocyclooctyne (DBCO)-488, and Western blot analysis.

DBCO labeling and immunoprecipitation were performed as follows. Prepared lysates were heated at 95°C for 10 minutes to denature proteins and increase access to internal AHA-bearing sequences. Iodoacetamide 10 mM, which decreases azide-independent labeling, was added to cooled lysates for 30 minutes at RT (17). Lysates were then immunoprecipitated using anti-IL-2Rα antibody directly conjugated magnetic beads. Conjugation was performed per manufacturer’s instructions (Click-&-Go, Click Chemistry Tools, Scottsdale, AZ). Following a 1h incubation at RT, beads (now containing bound protein) were washed and incubated with 5 μM DBCO-488 (Click Chemistry Tools) for 30 min at RT or overnight at 4°C. Beads were washed again and labeled protein was eluted with lithium dodecyl sulfate (LDS) containing sample buffer in preparation for SDS-PAGE.

Levels of DBCO-labeled IL-2Rα were normalized using either total protein (Revert™ 700 Total Protein Stain) or total IL-2Rα detected by either the anti-IL-2Rα antibody from BosterBio or anti-phospho-CD25 (ser268). The latter antibody was chosen for its strong signal and suggests that IL-2Rα is active. A linear regression analysis was then performed on normalized levels of labeled IL-2Rα and half-life was calculated via the following equation: t_1/2_= ln(2)/slope of decay (13).

### Statistics

Groups were compared using an unpaired, two-tailed t test with Welch’s correction. P values in figures are defined as follows: ns P > 0.05; * P ≤ 0.05; ** P ≤ 0.01; *** P ≤ 0.001; **** P ≤ 0.0001.

## Results

### IL-2Rα KO VSMC express IL-2Rα protein

To determine how IL-2Rα affects the function of VSMC, we isolated VSMC from IL-2Rα KO and WT mice. WT and KO mice were genotyped using the protocol provided by JAX^®^ (18). As a first step following isolation and culture, we examined expression of IL-2Rα protein in WT and KO VSMC by indirect immunofluorescence. VSMC isolated from WT mice expressed IL-2Rα primarily in the nucleus, consistent with our recent observations in human VSMC (Figure 1A, B). However, IL-2Rα was also detected in the nucleus of VSMC isolated from IL-2Rα KO mice. Similar results were obtained with antibodies from multiple sources. Although most antibodies detected IL-2Rα in the nucleus, two antibodies showed partial (Bioss) or predominantly (Genetex) membrane or cytoplasmic localization (Figure 1A and Supplemental Figure 1). Although we have not established the reason behind this differential detection, it likely relates to the fact that IL-2Rα is heavily glycosylated which may obscure some epitopes. Preadsorption with a known peptide epitope, available for the antibody obtained through BosterBio, abrogated staining (Figure 1C).

**Figure 1.**
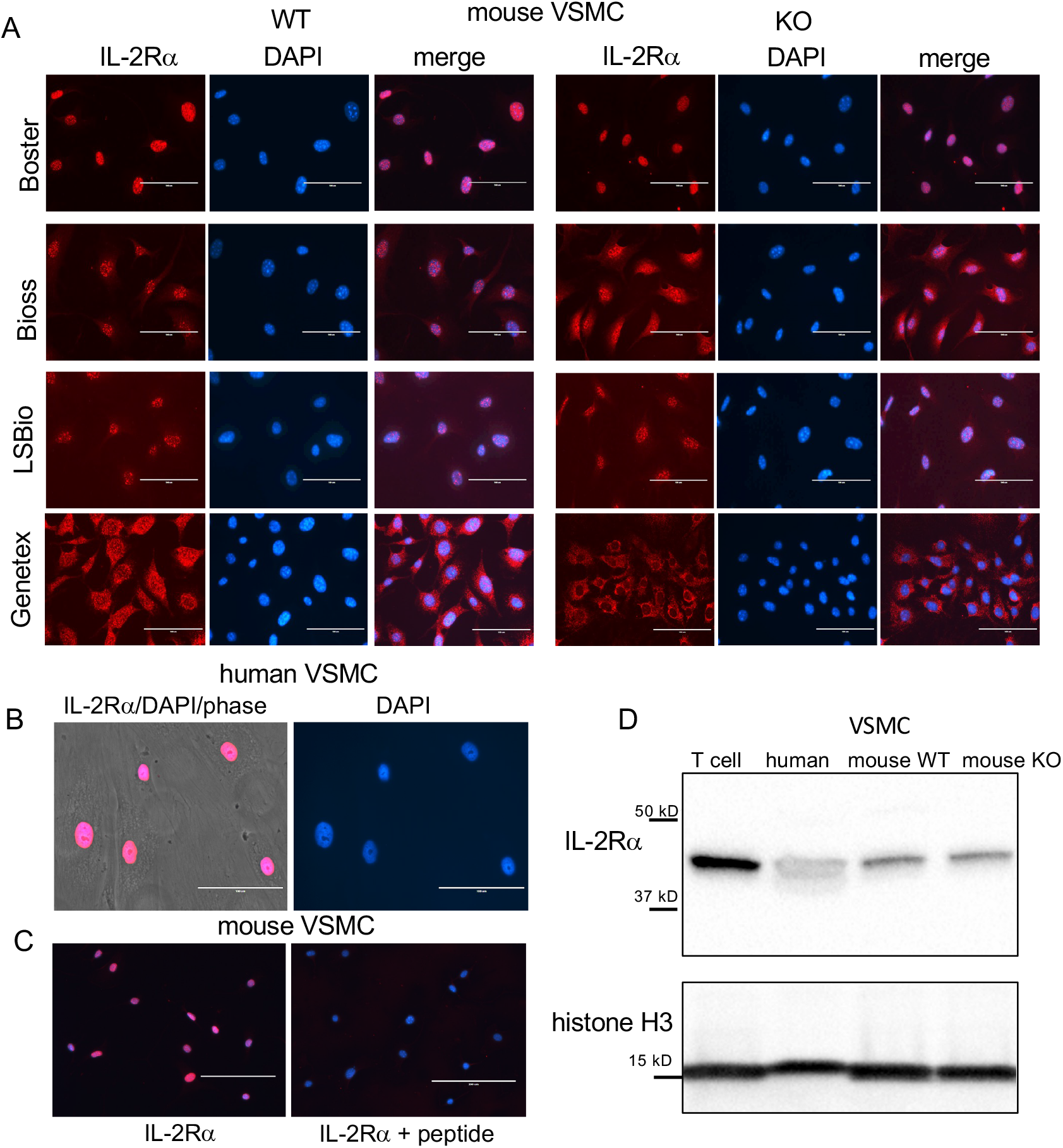
VSMC isolated from IL-2Rα KO mice express IL-2Rα protein. (A) VSMC, isolated from IL-2Rα WT and KO mice, were cultured to 60 - 70% confluency. After fixation and blocking, primary antibodies recognizing IL-2Rα, purchased from various sources as indicated, were applied to VSMC followed by the appropriate Cy5 conjugated secondary. Cells were imaged using an EVOS FL epifluorescence microscope (ThermoFisher Scientific). Scale bar = 100 μ. (B) Human VSMC were stained with the anti-IL-2Rα from BosterBio. Scale bar = 100 μ. (C) Murine VSMC were stained as in (A) using anti-IL-2Rα from Boster with/without pre-adsorption with immunizing peptide. Scale bar = 200 μ. (D) Lysates of VSMC or Jurkat T cells were separated by SDS-PAGE and analyzed for expression of IL-2Rα, using anti-IL-2Rα from Boster. Histone H3 was used as a loading control. Results shown are representative of >5 (A,B), 3 (C), and >5 (D) separate experiments.

IL-2Rα KO mice were generated by replacing exons 2 and 3 of IL-2Rα with a neomycin resistance gene (1). IL-2Rα is known to have multiple splice variants – three have been identified in mouse IL-2Rα and nine in human IL-2Rα (19). To ensure that the IL-2Rα protein detected was not produced by a splice variant, potentially excluding the neomycin resistance insert, we assessed the size of IL-2Rα protein produced by WT and KO VSMC using Western blot analysis. As seen in Figure 1D, WT and IL-2Rα KO VSMC expressed IL-2Rα of the same size, identical to that of human VSMC and Jurkat T cells (9). These results suggest that IL-2Rα KO VSMC express full-length IL-2Rα.

### IL-2Rα KO splenocytes express IL-2Rα protein

Because IL-2Rα expression and function are mainly studied in T cells, we asked whether lymphocytes from IL-2Rα KO mice express IL-2Rα protein. To this end, we isolated splenocytes from WT and IL-2Rα KO mice and examined IL-2Rα expression by flow cytometry in the presence or absence of stimulation with phytohemagglutinin (PHA) to upregulate membrane IL-2Rα. Since IL-2Rα appears mainly localized to the nucleus in VSMC, we also permeabilized cells to look for nuclear expression. To assess IL-2Rα expression, we used two of the same antibodies used in VSMC, sourced from Genetex and BosterBio (see Figure 1).

In non-permeabilized cells, a subset of WT splenocytes expressed IL-2Rα on their cell surface when stimulated with PHA (Figure 2A). A small subset of KO cells also expressed IL-2Rα. Upon permeabilization, however, IL-2Rα was detected by in the majority of WT splenocytes (70-80%). IL-2Rα was also detected in permeabilized KO splenocytes, however the frequency was lower than in WT cells, particularly those stained with the anti-IL-2Rα from Genetex. The intensity of staining was also decreased in permeabilized KO vs WT splenocytes (Figure 2B). Although not statistically significant, the frequency of permeabilized cells expressing nuclear IL-2Rα was decreased in WT cells stimulated with PHA vs those that were unstimulated. These results are consistent with our finding that, in VSMC, nuclear IL-2Rα is decreased in proliferating vs quiescent cells (unpublished observation).

**Figure 2.**
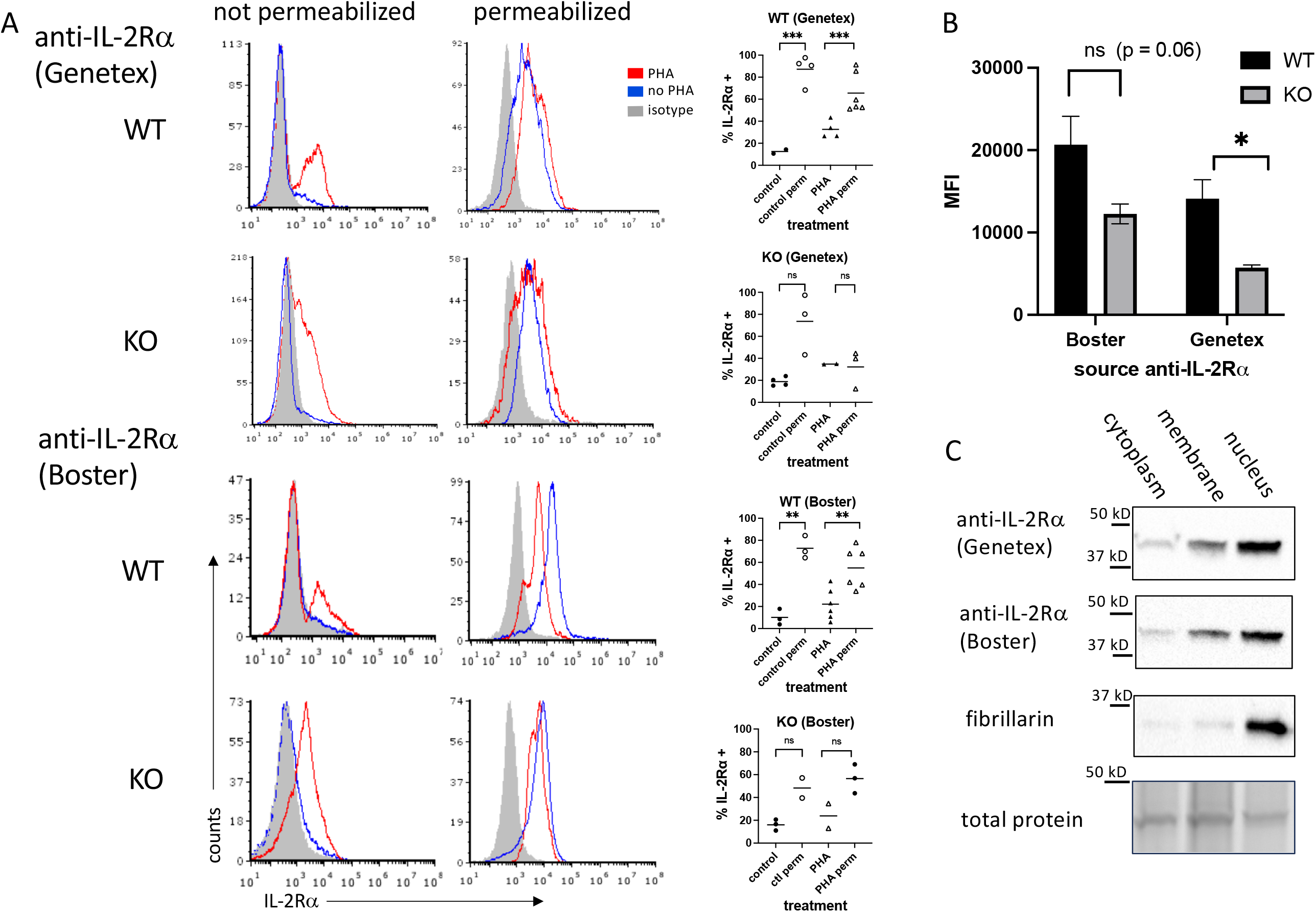
IL-2Rα is detectable in the nucleus of IL-2Rα WT and KO splenocytes. (A) Splenocytes, prepared fresh or following 48h of stimulation with PHA 1 μg/ml, were stained with anti-IL-2Rα antibodies or isotype controls as indicated with/without first permeabilizing via methanol. Graphs to the right of histograms represent a summary of individual values from multiple experiments. (B) represents the average MFIs ± SEM, derived from (A), of permeabilized splenocytes stained with anti-IL-2Rα antibodies as indicated. (C) Splenocytes were stimulated with PHA for 48h, pelleted and fractionated into membrane, cytoplasm, and nuclear fractions, then analyzed by Western blot probing with antibodies as indicated. Blots represent lysates processed in parallel. Total protein, obtained from stain-free gels (Bio-Rad), shows the single most prominent band present across all samples. Results shown are representative of ≥ 3 experiments.

These results suggested that IL-2Rα could be localized to the nucleus in lymphocytes. To address this question, we separated splenocyte lysates into membrane, cytoplasm, and nuclear fractions. Consistent with our findings in VSMC, IL-2Rα was present in the nucleus (Figure 2C).

We then asked whether a commonly used anti-IL-2Rα antibody, rat anti-mouse IL-2Rα clone PC61.5, would yield similar results (20–23). Non-permeabilized WT splenocytes stimulated with PHA yielded strong expression, as expected, and similarly treated KO cells were negative (Figure 3A). However, permeabilization yielded only low levels of IL-2Rα in WT cells and KO cells were negative. Given the nuclear localization of IL-2Rα (Figures 1,2), we reasoned that a potential association with DNA might be blocking some epitopes. To test this question, we digested permeabilized cells with Benzonase^®^, a genetically engineered endonuclease that degrades all forms of DNA and RNA. IL-2Rα was strongly detected in permeabilized WT and KO splenocytes following Benzonase^®^ digestion, although the level of expression per cell (mean channel fluorescence) was much higher in WT cells (Figure 3B). Pre-adsorption of PC61.5 with purified mouse IL-2Rα protein abrogated this staining (Figure 3A).

**Figure 3.**
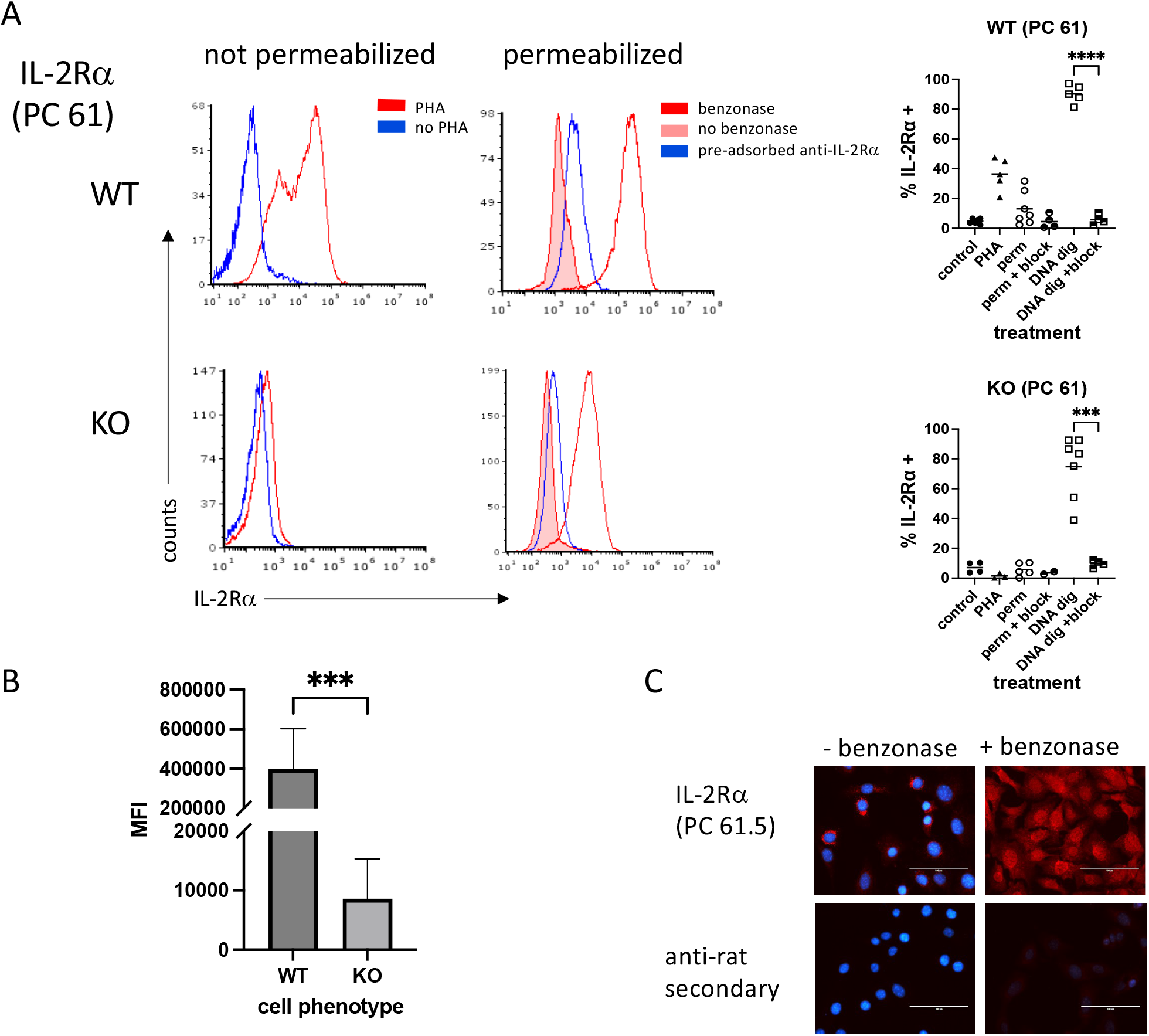
IL-2Rα is detected in permeabilized splenocytes by rat anti-mouse IL-2Rα (clone PC 61) following DNA digestion. (A) Splenocytes were prepared as described in Figure 2. A subset of methanol permeabilized cells were treated with Benzonase® 250U/10^6^ cells or enzyme buffer for 50 minutes at 37°C prior to staining. In a subset of samples, anti-IL-2Rα/clone PC 61 was pre-adsorbed with an excess of purified mouse IL-2Rα. Graphs to the right of histograms represent a summary of individual values from multiple experiments. “Block” indicates use of pre-adsorbed anti-IL-2Rα. (B) represents the average MFIs ± SEM, derived from (A), of permeabilized splenocytes treated with Benzonase®. (C) VSMC were permeabilized with methanol and treated with Benzonase® or buffer, as indicated above, then stained with anti-IL-2Rα/clone PC 61. Note that digestion with Benzonase® eliminates DAPI staining as expected. Scale bar = 100 μ.

Although prior attempts at using clone PC61.5 to detect IL-2Rα in VSMC yielded poor results, based on the above findings we digested methanol-fixed VSMC with Benzonase^®^. Digestion of mouse VSMC with Benzonase^®^ allowed for the detection of IL-2Rα in the nucleus, consistent with our findings in T cells (Figure 3C). In addition, these results suggest that at least a portion of nuclear IL-2Rα binds DNA.

### Genomic sequencing demonstrates IL-2RA deletion

In light of our results demonstrating that both IL-2Rα KO splenocytes and VSMC produce IL-2Rα, we sought to establish its source. As a first step, we asked if the deletion in the *IL2RA* gene was intact. To this end, we sequenced genomic DNA isolated from WT and KO VSMC. Targeted genome sequencing of the *IL2RA* gene from KO cells revealed correct deletion of exons 2 and 3 as initially reported by Willerford, et al (1) (Figure 4).

**Figure 4.**
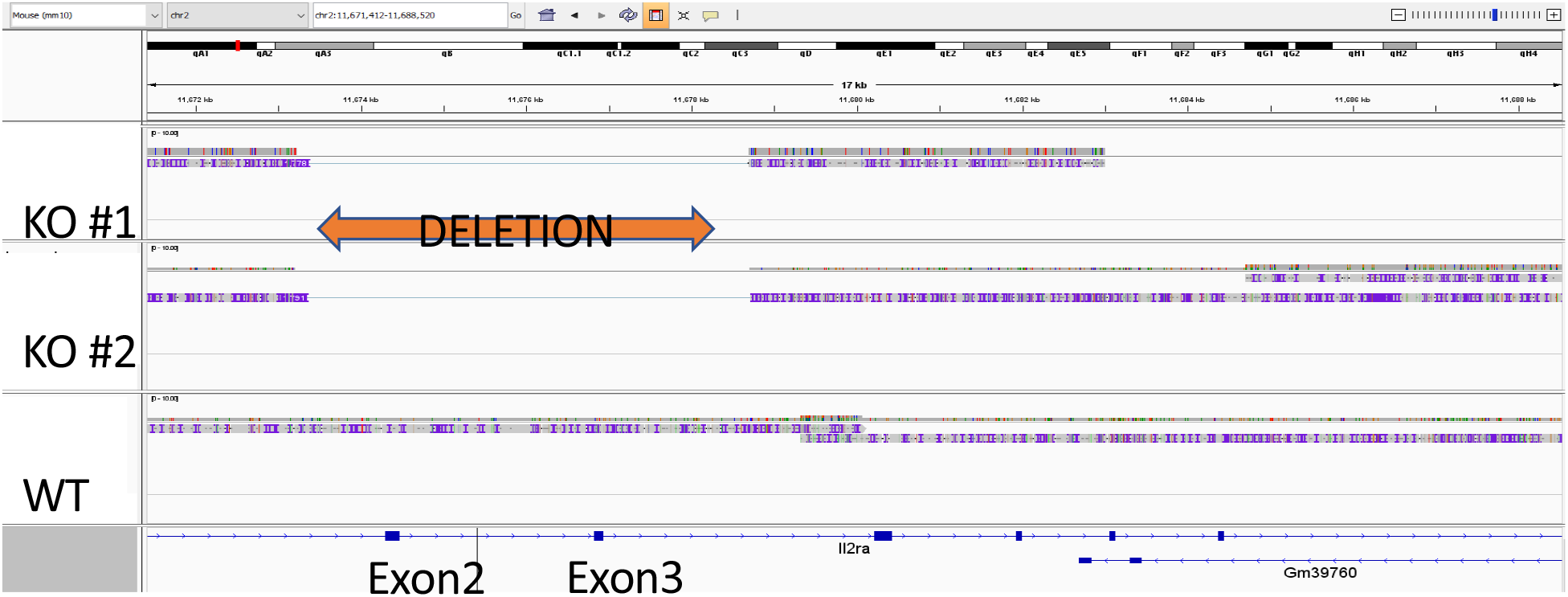
Exons 2 and 3 of IL-2Rα gene are deleted in IL-2Rα KO mice. Genomic DNA isolated from WT and KO VSMC underwent targeted sequencing, with a focus on the IL-2Rα locus. Exons 2 and 3 of IL-2Rα are deleted as reported by Willerford, et al (1).

### Multiple tissues in IL-2Rα KO mice have low levels of the IL-2RA gene

Given that *IL2RA* was correctly deleted, we considered other sources of the WT gene. One potential source of the WT *IL2RA* gene is maternal transfer. Maternal microchimerism, the transfer of cells from mother to fetus, is well established in humans and mice wherein maternal DNA/cells can persist for decades in offspring (24–28). Because IL-2Rα KO mice are infertile, they are generated by mating IL-2Rα heterozygotes; consequently, heterozygous maternal cells are likely present in IL-2Rα KO mice. Maternal microchimerism may therefore represent result a potential source of IL-2Rα in KO offspring of heterozygous females. Our laboratory previously demonstrated that maternal microchimerism was responsible for the presence of IL-2 in IL-2 KO mice, which obscured severe defects in thymic development (27).

Using the *IL2RA* gene as an indicator of maternal microchimerism, we extracted genomic DNA from IL-2Rα KO tissues and looked for evidence of *IL2RA*, using primers that amplify a segment within exon 2 that should be absent in KO mice (1). Relative levels of *IL2RA* were assessed by qPCR of genomic DNA. QPCR of maternal markers in genomic DNA of offspring has been used by other investigators to detect maternal microchimerism (24). As seen in Figure 5A, 6/6 spleens, 6/6 kidneys, and 3/3 hearts from adult KO mice had detectable (RQ > 0 with one of the KO spleens as reference) levels of *IL2RA*. As further evidence for the presence of maternal cells in offspring of IL-2Rα het breeders, we assessed spleens from IL-2Rα WT offspring of heterozygous parents for the neomycin resistance gene used in the generation of IL-2Rα KO mice. Five of five spleens tested had detectable levels of the *NeoR* gene (Figure 5B). As expected, control spleens from unrelated mice that were not offspring of IL-2Rα het mice had no detectable *NeoR* gene.

**Figure 5.**
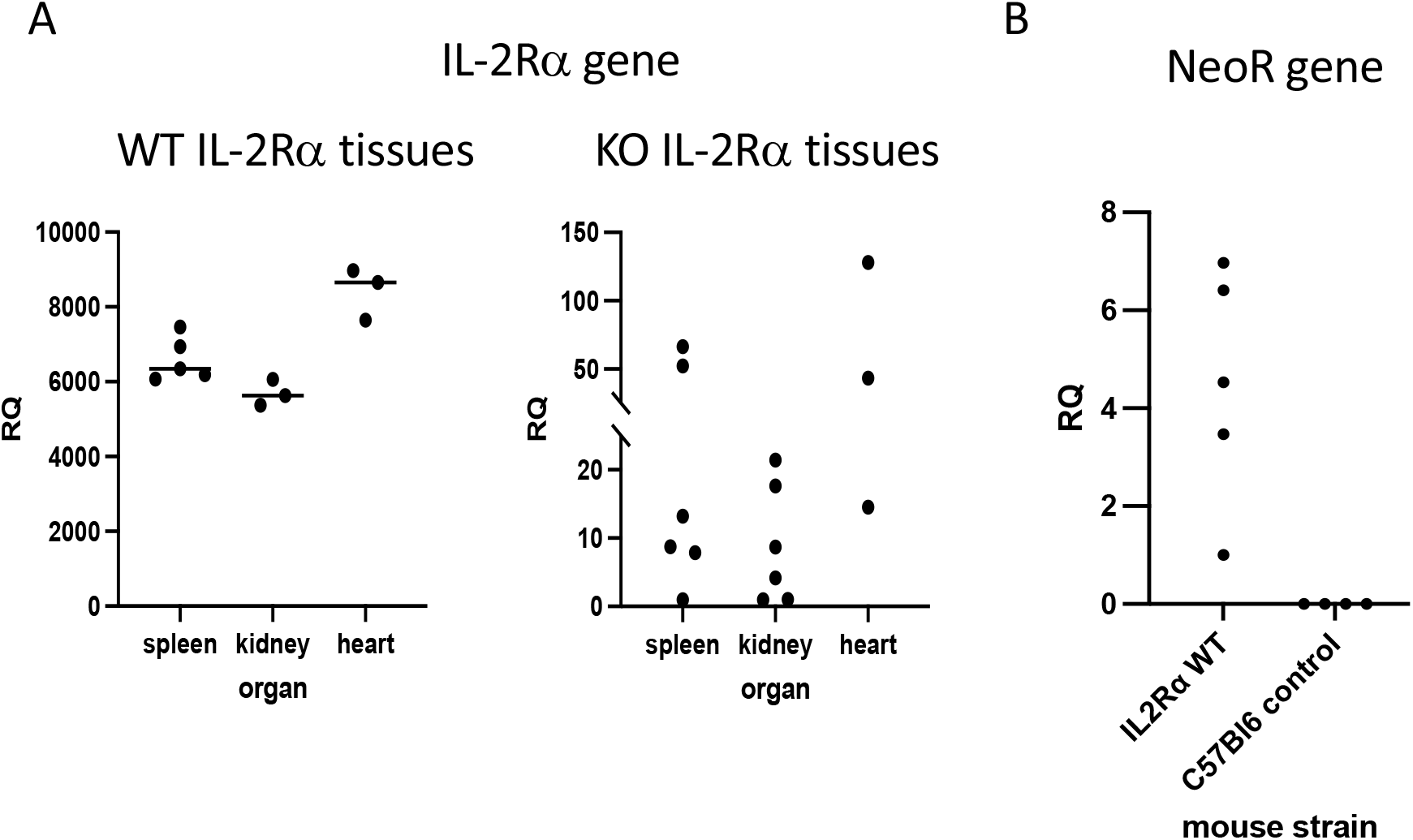
IL-2Rα gene is present in IL-2Rα KO tissues. Genomic DNA was isolated from various organs as indicated. Levels of *IL2RA* (A) or *Neo* resistance (B) DNA were measured in each tissue by qPCR. The relative quantity (RQ) of IL-2Rα or NeoR DNA was measured relative to a single KO spleen (A) or a WT spleen (B).

### G418 decreases IL-2Rα protein expression in IL-2Rα KO VSMC

As previously mentioned, IL-2Rα KO mice were generated by replacing exons 2 and 3 of IL-2Rα with a neomycin resistance gene insert as a selectable marker (1). If IL-2Rα KO VSMC contained low numbers of IL-2Rα heterozygous cells, as our results suggest, then treatment with neomycin might eliminate these cells if they are less resistant to neomycin than KO cells. In turn, IL-2Rα protein expression should decrease. To test this hypothesis, we cultured WT and KO VSMC with increasing concentrations of G418 (a neomycin analog). G418 nearly eliminated WT cells at a concentration of 2 mg/ml (Figure 6A). However, many VSMC isolated from KO mice survived this treatment. IL-2Rα production by these cells following treatment with G418 was significantly diminished compared to untreated VSMC isolated from IL-2Rα KO mice as shown by both immunofluorescence and Western blot analysis (Figure 6A, C). These data suggest that the neomycin resistance protein is expressed and functional in IL-2Rα KO VSMC, and that treatment of KO VSMC with G418 decreased IL-2Rα protein originating from heterozygous cells.

**Figure 6.**
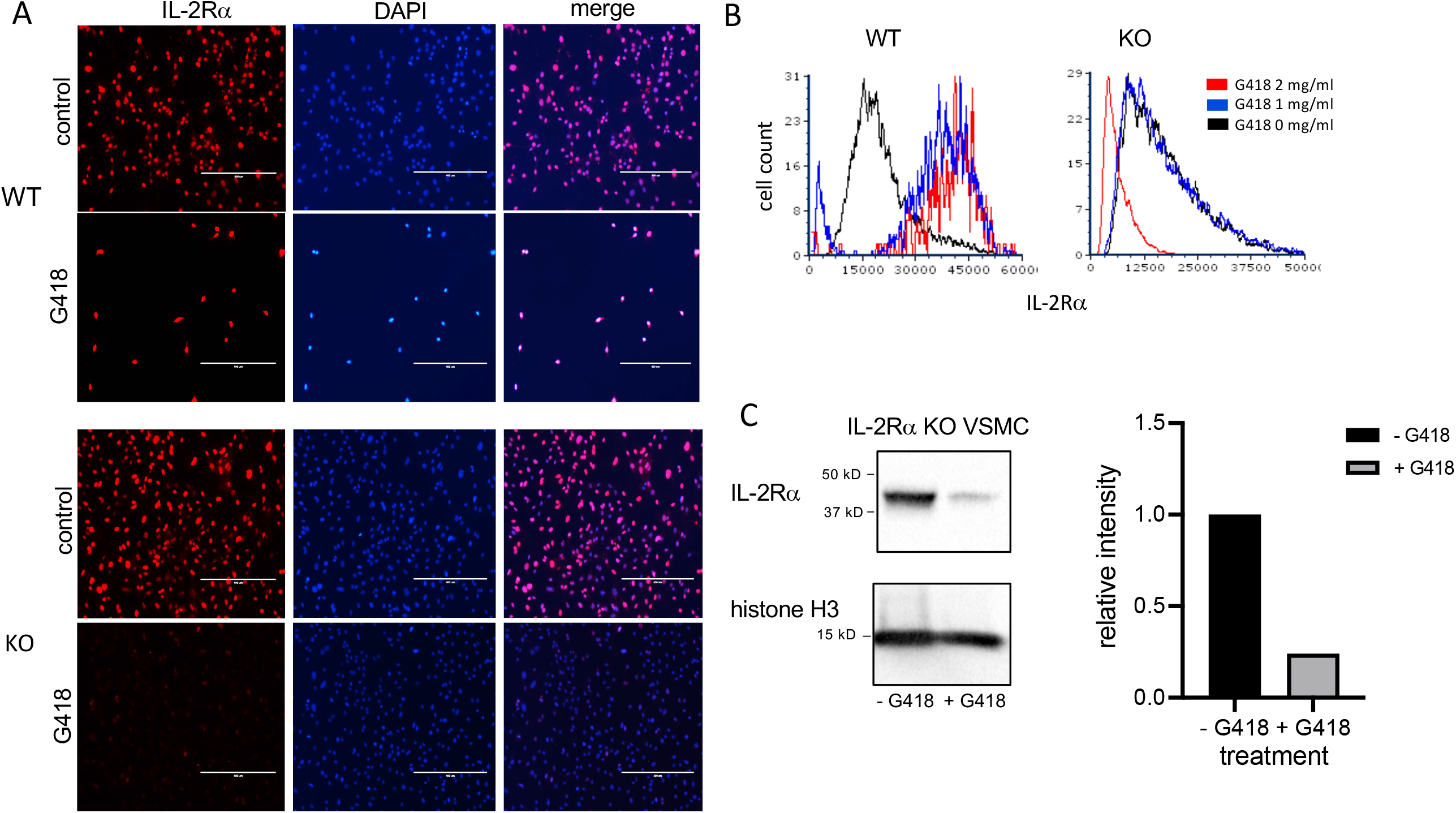
G418 decreases IL-2Rα protein expression in VSMC isolated from IL-2Rα KO mice. (A) VSMC, isolated from IL-2Rα WT and KO mice, were cultured in the presence/absence of G418 at 2 mg/ml. Cells were probed for IL-2Rα expression using the anti-IL-2Rα from Boster as described in Figure 1. Scale bar = 400 μ. (B) The intensity of IL-2Rα staining in all cells from (A) was measured in an image cytometer and converted to a histogram using flow cytometry software. (C) KO VSMC, cultured as described in A, were lysed. Extracted proteins were separated by SDS-PAGE and analyzed by Western blot for expression of IL-2Rα using the anti-IL-2Rα antibody from Boster. Intensity of the IL-2Rα band from each treatment was expressed as a ratio of IL-2Rα/histone H3. Densitometry was performed using Image Lab software from Bio-Rad Laboratories. Results shown for A,C are representative of >3,2 separate experiments respectively.

### IL-2Rα protein is transferred between cells

Although our first attempt at clearance of IL-2Rα protein by treatment with G418 significantly decreased IL-2Rα protein expression, we were unable to eliminate IL-2Rα despite ongoing treatment with G418 and/or increased dosage (Figure 7). Upon performing formal kill curves with G418, we observed that the difference in resistance between heterozygous and homozygous cells is probably not large enough to differentially eliminate all heterozygous cells (Supplemental Figure 2).

**Figure 7.**
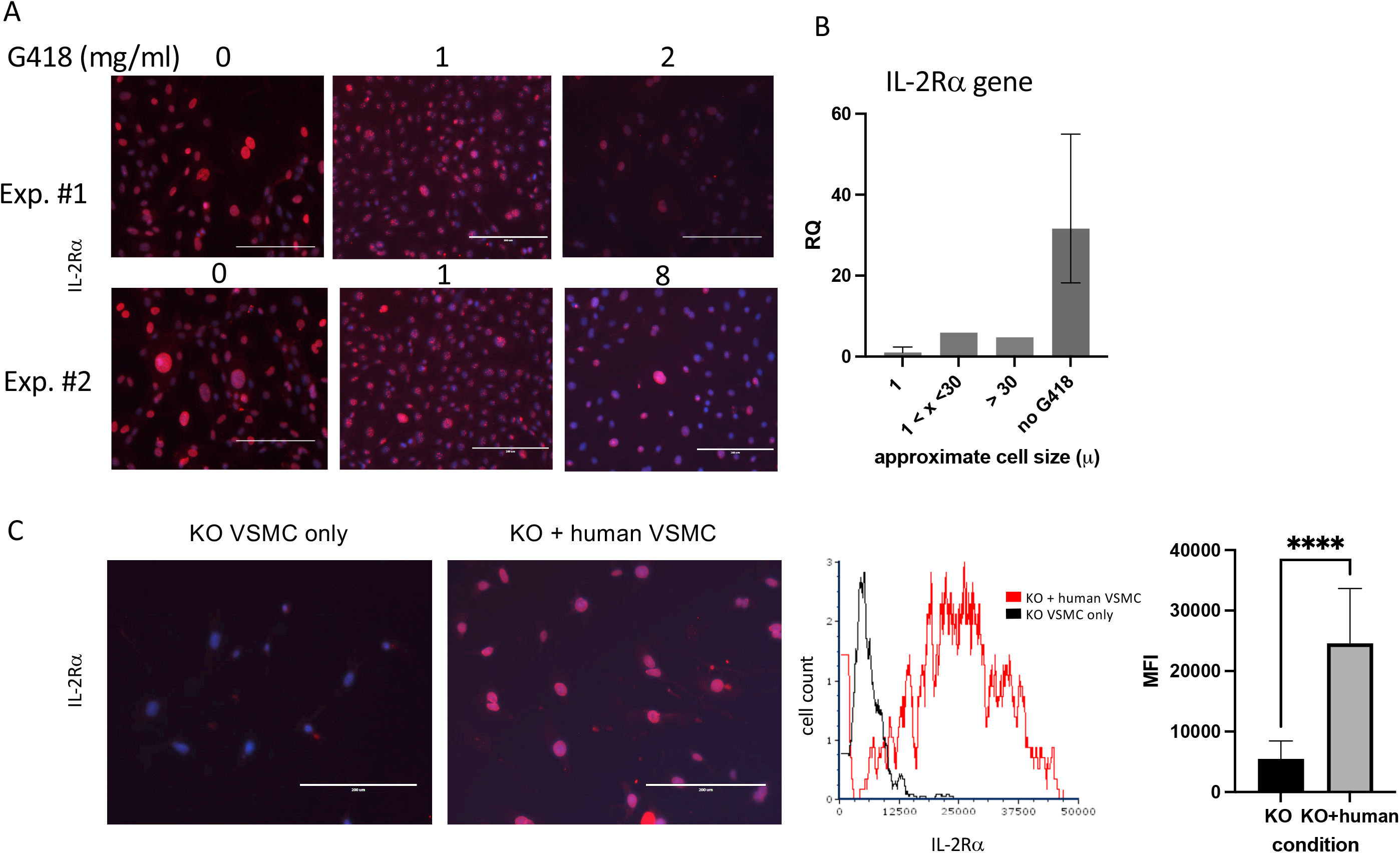
IL-2Rα is transferred between cells. (A) VSMC were cultured for 96h with increasing concentrations of G418 as indicated. IL-2Rα protein expression was detected using the anti-IL-2Rα antibody from BosterBio. (B) VSMC, cultured in G418, were separated by size using 1 and 30 micron filters as indicated. Genomic DNA was isolated from cell pellets and levels of *IL2RA* DNA were quantified relative to the 1 micron cells. (C) Human VSMC were placed in transwell inserts and co-cultured with murine VSMC for 96h and compared to murine VSMC without co-culture. IL-2Rα protein expression was detected using the anti-IL-2Rα antibody from BosterBio. Intensity of IL-2Rα staining in all cells was measured in an image cytometer (Cytation) then converted to a histogram or bar graph using flow cytometry or data analysis software. Scale bar = 200 μ.

As part of efforts to identify the source of ongoing IL-2Rα production, we noted that VSMC were heterogeneous in size, and that large cells appeared to express high levels of IL-2Rα protein (Figure 7A). Using filters (Pluriselect, Leipzig, Germany), we separated cells into those that passed through a 1 micron filter from the remainder that did or did not pass through a 30 micron filter. We then compared levels of *IL2RA* DNA amongst these cells and with those that had not received G418. As seen in Figure 7B, larger cells contained relatively more *IL2RA* DNA than smaller cells, and those without G418 treatment contained the most *IL2RA* DNA, consistent with our observations of IL-2Rα protein expression (Figures 6,7).

Given the above findings, combined with our observation that most VSMC isolated from IL-2Rα KO mice expressed at least low levels of IL-2Rα protein, we asked whether IL-2Rα could be transferred between cells. To this end, we co-cultured IL-2Rα KO VSMC with human VSMC using a transwell insert, in which human VSMC were suspended over KO VSMC and separated by a filter. We chose human VSMC for their robust IL-2Rα protein expression (Figure 1). IL-2Rα was easily detectable in IL-2Rα KO VSMC that had been co-cultured with human VSMC, compared to those that were not (Figure 7C). These results suggest that IL-2Rα protein is transferred between cells by a means not requiring cell contact, likely through extracellular vesicles such as exosomes or apoptotic bodies.

### Half-life of IL-2Rα

While the previous results suggest that IL-2Rα protein present in KO mice originated from maternal microchimerism, the low levels of *IL2RA* DNA detected did not seem consistent with the comparatively high levels of protein. One way in which protein levels may be regulated is through their half-life. In a recent report by Chen, et al, a systematic study of half-lives of proteins in human HepG2 cells showed that most proteins had half-lives ranging from 4-14 hours (29).

Given the easily detectable IL-2Rα protein evident in KO mice, we hypothesized that the half-life of IL-2Rα is long, likely days versus hours. To measure the half-life of IL-2Rα, we performed a pulse chase experiment, in which we labeled newly synthesized proteins with a methionine analog, AHA, then assessed their degradation over time through loss of labeled proteins. Incorporated AHA was detected by fluorescently labeled DBCO, which binds the azido group in AHA through strain promoted alkyne-azide cycloaddition, a copper-free click reaction (14). Based on our results showing that IL-2Rα is localized primarily to the nucleus, we focused on the nuclear fraction. IL-2Rα protein was isolated by immunoprecipitation using the anti-IL-2Rα from BosterBio, and then detected with either the anti-IL-2Rα from BosterBio or an anti-IL-2Rα (phospho-ser 268). Levels of DBCO-labeled IL-2Rα, assessed by densitometry, were normalized based on either total protein or total IL-2Rα protein per band. Normalizing with total IL-2Rα protein was sometimes difficult since the DBCO label variably impacted recognition of IL-2Rα (Supplemental Figure 3). Based on 3 separate experiments, the half-life of IL-2Rα was 8.4 ± 2.5 days (Figure 8). These results demonstrate that the half-life of IL-2Rα is longer than most cellular proteins and may explain the discrepancy between levels of *IL2RA* DNA and protein observed in IL-2Rα KO mice.

**Figure 8.**
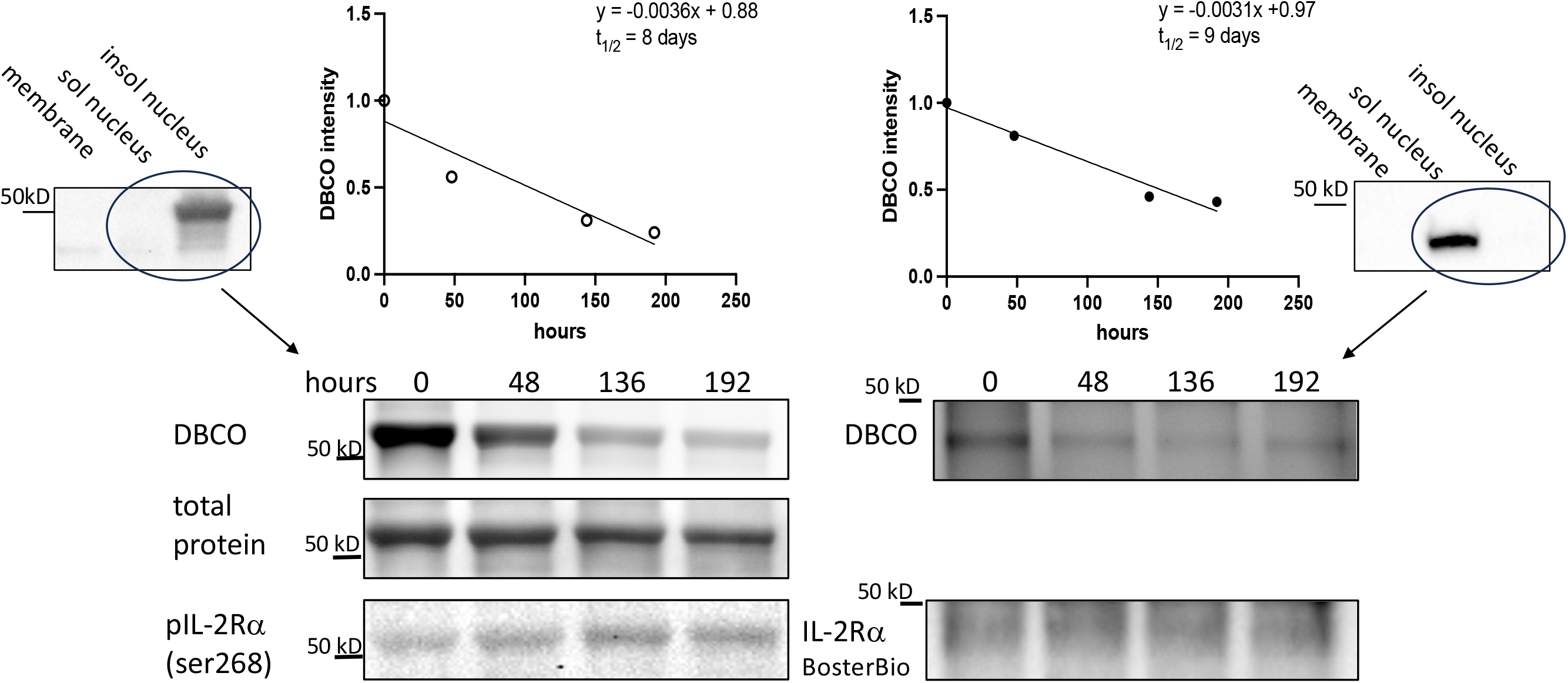
IL-2Rα exhibits a prolonged half-life. VSMC were labeled with AHA-containing, methionine free media for 48h then chased with complete media for the durations above. IL-2Rα was then isolated by immunoprecipitation with anti-IL-2Rα from BosterBio. AHA-labeled IL-2Rα was detected by DBCO-488 and total IL-2Rα was detected using anti-IL-2Rα, anti-phospho IL-2Rα ser268, or total protein stain as indicated. DBCO intensities, normalized to total IL-2Rα, were graphed on a linear scale and linear regression analysis was performed to determine the protein half-life. Blots at the left and right edges of the figure show VSMC fractionated into membrane, soluble nuclear, and insoluble nuclear fractions (both nuclear fractions digested with Benzonase^®^) and probed with antibodies as indicated. Circled lanes indicate the fractions that were combined for immunoprecipitation. The half-life on the left hand side of the figure was calculated using the band detected by anti-phospho IL-2Rα ser268 and total protein (of that band) for normalization. The half-life on the right hand side was calculated using the band detected by anti-IL-2Rα from BosterBio and immunodetection for normalization. In the above experiments, immunoprecipitated IL-2Rα was eluted using a denaturing buffer. Use of a non-denaturing buffer yielded a similar half-life of 8.25 days.

### IL-2Ra KO VSMC

Responses of IL-2Rα KO VSMC to sera, evident in routine culture, began to give us insight into the differences between WT and KO VSMC. Doubling times were significantly shorter in KO vs WT VSMC (Figure 9). KO VSMC were also much smaller than WT and took up less DAPI; these differences were heightened upon stimulation with FBS (Figure 9). The latter observations are consistent with the finding that DNA content is directly proportional to cell size (30). In comparing nuclear area and DNA content of WT and KO VSMC, however, we noted that the decrease in DNA content was out of proportion to the decrease in nuclear area. KO nuclei were approximately half the size of WT, but only had 1/6 the amount of DNA. This finding suggests that KO VSMC are hypodiploid, which implies a severe dysregulation of cell proliferation and/or division. Hypodiploid cells are mainly associated with an aggressive form of acute lymphoblastic leukemia (31,32).

**Figure 9.**
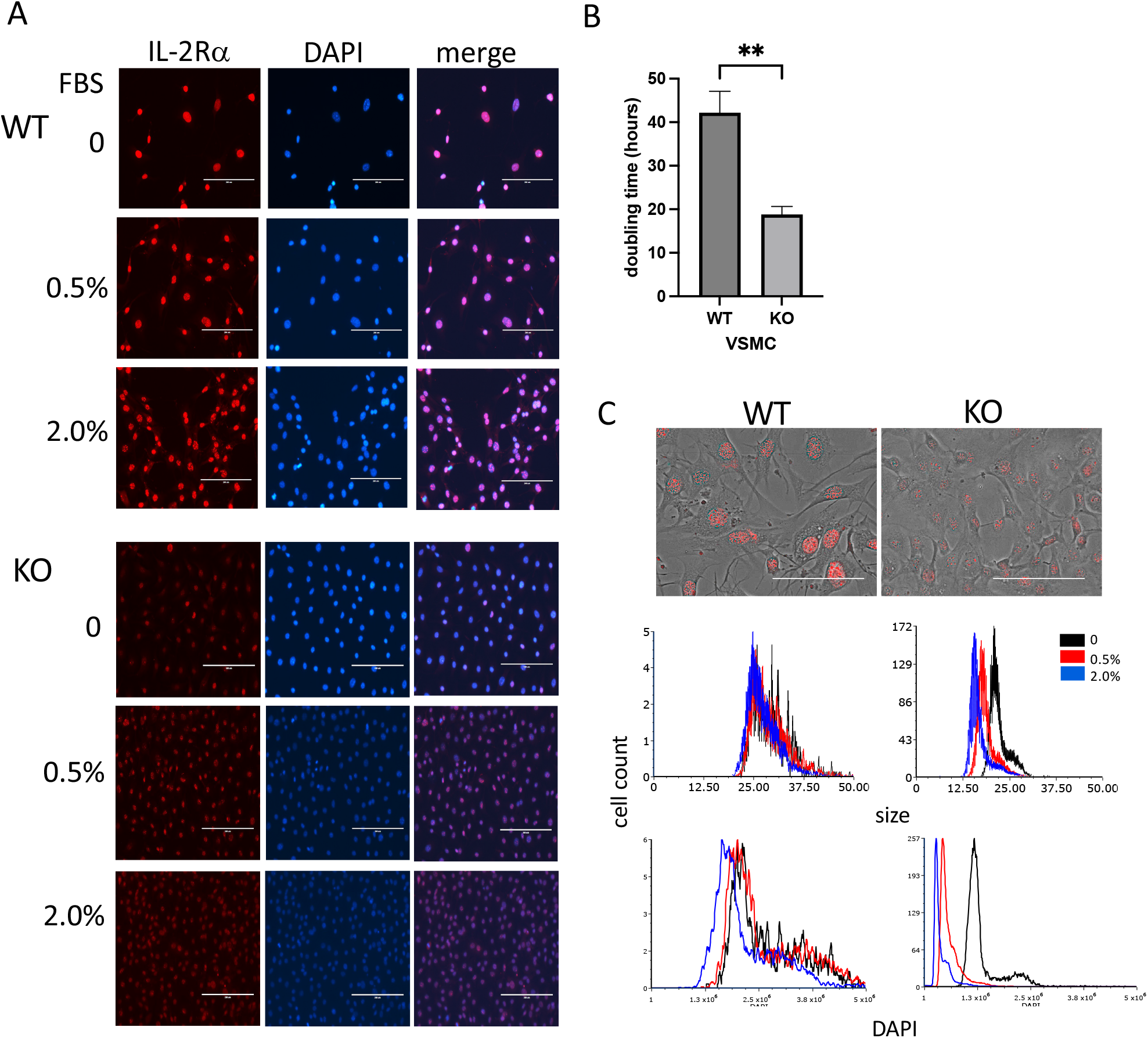
IL-2Rα KO VSMC are small and hypodiploid. (A) VSMC, isolated from IL-2Rα WT and KO mice, were cultured for 72h in serum free media with increasing concentrations of FBS. Cells were probed for IL-2Rα expression using the anti-IL-2Rα from Boster as described in Figure 1. Scale bar = 200 μ. (B) Doubling times of WT and KO VSMC were calculated using three time points over 72h (41). Data represent average ± SEM of 5 separate experiments. (C, top) Phase and IL-2Rα overlay images from WT and KO VSMC in serum free media and probed for IL-2Rα as in (A). Scale bar = 100 μ. (C, middle and bottom) Nuclei of VSMC from (A) were imaged for area and DAPI intensity using a Cytation imaging plate reader (Biotek). Data accrued from these images were then processed using flow cytometry software. Results shown are representative of 3(A,C) separate experiments or 5 (B) collated experiments.

## Discussion

Knock out mice are often used to determine the function of a protein. Both IL-2 and IL-2Rα KO mice were instrumental in determining that the IL-2/IL-2R system plays a major regulatory role in the immune system by promoting the survival of T regulatory cells and through activation-induced cell death (1,3,33). Our results show that IL-2Rα is diminished, but not absent, in IL-2Rα KO mice. Furthermore, our data suggest that the source of IL-2Rα is maternal microchimerism. The infertility of IL-2Rα KO mice, mandating a heterozygous mother, combined with the long half-life of IL-2Rα, creates a unique situation in which substantial amounts of IL-2Rα are present in KO mice. The predominantly nuclear localization of IL-2Rα, reported here, is likely responsible for its lack of detection before now.

Our studies of IL-2Rα KO mice and the mechanisms contributing to its incomplete elimination have led to several remarkable discoveries including: (1) IL-2Rα is localized primarily to the nucleus, (2) IL-2Rα protein has a long half-life, and (3) IL-2Rα is transmitted between cells. First, and foremost, is our observation that IL-2Rα is localized primarily to the nucleus. Prior reports addressing the potential nuclear localization of IL-2R subunits are rare. In 1988, Jothy, et al showed that IL-2Rα was transiently localized to the nucleus of concanavalin A supernatant-stimulated HT-2 cells (34). Conversely, Fujii, et al, showed that neither IL-2Rβ nor IL-15Rα could be transported to the nucleus, however they did not test IL-2Rα (35). Our report adds to this literature, clearly demonstrating that IL-2Rα localizes to the nucleus. In this location, our data show that IL-2Rα deficient cells become smaller and lose DNA when stimulated to proliferate. These results are consistent with our finding that quiescent and senescent cells have increased nuclear IL-2Rα relative to proliferating cells (unpublished observation). While not formally studied, our methods also intimate that IL-2Rα binds DNA, given that we were unable to immunoprecipitate IL-2Rα without first digesting DNA. Taken together, these findings suggest that that IL-2Rα contributes to the regulation of cell division and/or acts as a transcription factor.

In addition to its nuclear localization, we have also determined that IL-2Rα has a long half-life relative to most proteins. Systemic studies of protein half lives in cells and tissues have shown that mitochondrial and nuclear proteins have the longest half-lives, consistent with the predominantly nuclear localization of IL-2Rα reported here (29,36). Proteins with long half-lives tend to play key structural or functional roles within cells. Histones, for example, with half-lives ranging from 15 days in dividing cells to 240+ in non-dividing cells, contribute to the organization of chromatin which in turn regulates cell division, DNA damage responses, gene expression, and cell fate (29,37,38). Several mitochondrial proteins have half-lives on the order of 7 days; these longer-lived proteins were found to contribute to the stability of the electron transport chain (39). The long half-life of IL- 2Rα, demonstrated here, provides additional support to the concept that IL-2Rα has a newfound, fundamental role in regulating cell size and proliferation.

Finally, our investigation into the incomplete elimination of IL-2Rα mice revealed that IL-2Rα is shared/transported between cells. Given that cell contact was not required for this transmission, it is likely that IL-2Rα was transported through extracellular vesicles such as an exosomes or apoptotic bodies. Extracellular vesicles are a major means of intercellular communication and have been shown to participate in many biological processes including differentiation, proliferation, apoptosis, motility, and others (40). If IL-2Rα plays a cell intrinsic role in regulating proliferation, as suggested by our findings, then transmission of IL-2Rα by its “producers” may promote the normal function of surrounding KO cells and attenuate the consequences of IL-2Rα deficiency.

As previously described, we used G418 resistance to decrease the production of IL-2Rα, originating from IL-2Rα heterozygous cells, in VSMC isolated from IL-2Rα KO mice. This method was partially successful (Figures 6,7); however, to date we have not been able to completely eradicate IL-2Rα. Reasons for this ongoing incomplete eradication likely include (1) insufficient difference between G418 resistance of homozygous vs heterozygous cells, (2) lack of means to detect and exclude live IL-2Rα producing cells, and (3) techniques such as clonal dilution are challenging in primary VSMC as their viability at the single cell level is poor. Our laboratory is in the process of finding other methods of detecting and eliminating IL-2Rα producing cells to establish a complete IL-2Rα KO phenotype both *in vitro* and *in vivo*.

In summary, our data reveals the surprising finding that IL-2Rα KO mice express significant amounts of IL-2Rα protein. Our initial assessment of IL-2Rα KO VSMC suggests that IL-2Rα, localized to the nucleus and independent of IL-2, plays an unexpected role in the regulation of cell proliferation.

## Author Contributions

VW, KD, and GC performed experiments, analyzed data and contributed to experimental design. LW designed experiments, analyzed data, and wrote the manuscript. All authors critically reviewed and edited the manuscript.

## Acknowledgements

The authors would like to extend a heartfelt thanks to the Microarray and Microbiome Sequencing Core Facility at University of Texas Southwestern Medical Center for performing the quantitative PCR experiments and targeted genomic sequencing assay for the present study.

