## Supplemental figures for "IL-2Rα KO mice exhibit maternal microchimerism and reveal nuclear localization of IL-2Rα in lymphoid and non-lymphoid cells"

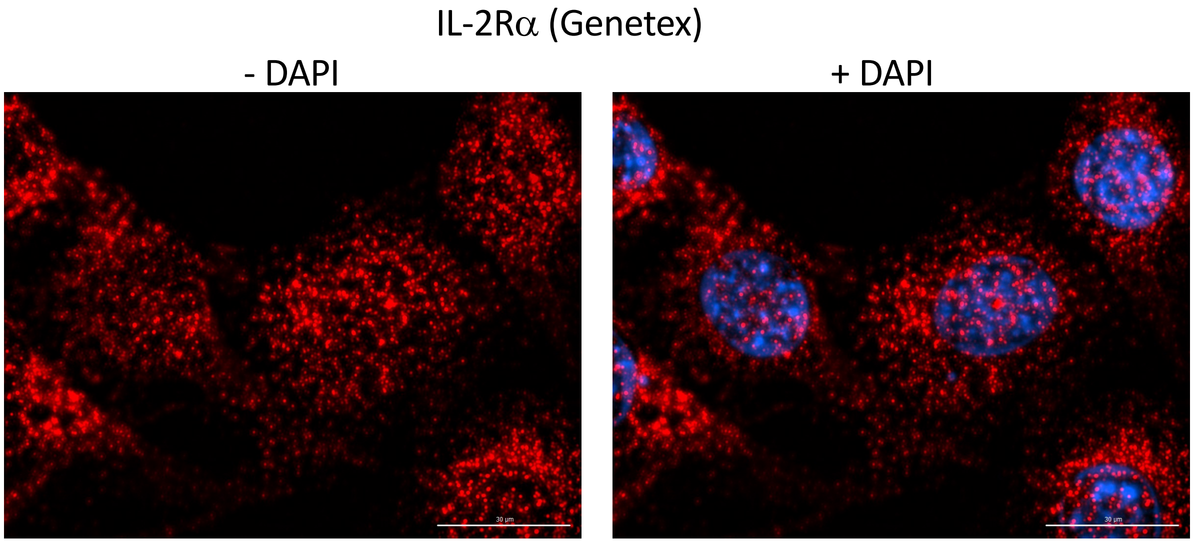


**Supplemental Figure 1. Mouse anti-IL-2Rα, clone 1B5D12, detects IL-2Rα in both the nucleus and cytoplasm/membrane.** Mouse VSMC were grown to approximately 70% confluence, then fixed and stained using a mouse monoclonal anti-IL-2Rα antibody, clone 1B5D12. Images were captured as a Z stack with a Cytation multi-mode imaging plate reader using a 1.6 μ depth of field. The image shown is 1.6 μ into the stack. Scale bar = 30 μ.

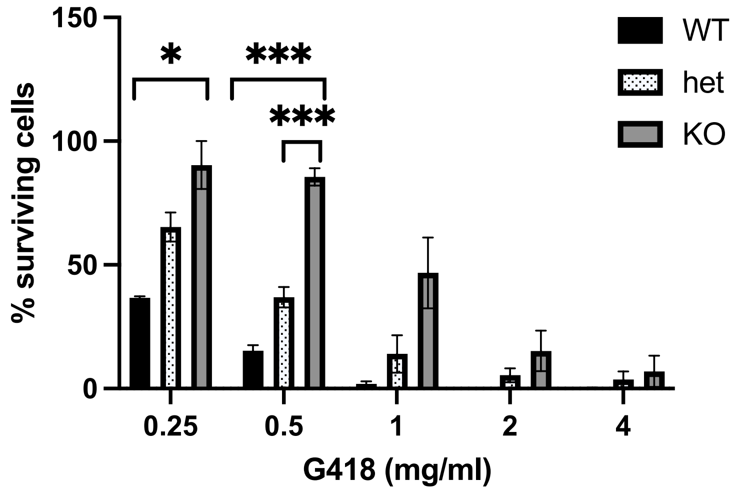


**Supplemental Figure 2. Sensitivity of WT, heterozygous, and IL-2Rα KO VSMC to G418**. VSMC, isolated from mice with genotypes as indicated, were cultured with increasing concentrations of G418 for 96h. Remaining cells were counted using a nuclear stain. Survival is expressed as a percentage of remaining cells using cells cultured without G418 as the denominator. Results shown represent the average ± SEM of three experiments.

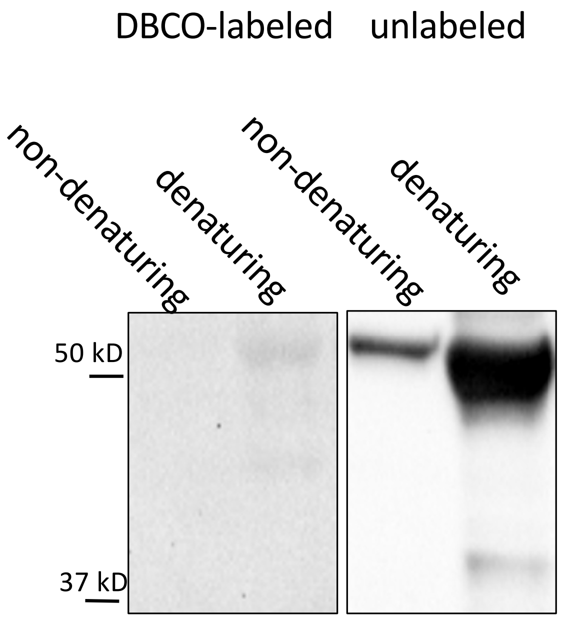


**Supplemental Figure 3. DBCO label interferes with detection of IL-2R$\alpha$ by Western blot.** IL-2R$\alpha$ was isolated from nuclear fractions of VSMC lysates by immunoprecipitation using a rabbit polyclonal anti-IL-2R$\alpha$ from BosterBio. IL-2R$\alpha$ protein was eluted from beads (directly conjugated with antibody) using either a non-denaturing or denaturing buffer. DBCO was added to a portion of the lysates as indicated. Lysates were subjected to Western blot analysis and probed with anti-phospho IL-2R$\alpha$ (ser268). Similar issues with detection were noted when probing with rabbit anti-IL-2R$\alpha$ polyclonal (BosterBio) or mouse anti-IL-2R$\alpha$ monoclonal (clone 1B5D12, Genetex) antibodies.
